# Laboratory evolution of multiple *E. coli* strains reveals unifying principles of adaptation but diversity in driving genotypes

**DOI:** 10.1101/2020.05.19.104992

**Authors:** Erol S. Kavvas, Maciek Antoniewicz, Christopher Long, Yang Ding, Jonathan M. Monk, Bernhard O. Palsson, Adam M. Feist

## Abstract

Fitness landscapes are a central concept in evolutionary biology and have been thoroughly detailed in terms of genotypes. However, our understanding of the selected metabolic and gene expression adaptations, and their dependence on genetic background, remains limited. Here, we reveal multi-scale adaptation principles in the *E. coli* species by taking multi-omics measurements of six different strains throughout their adaptive evolution to glucose minimal media. Statistics and matrix factorization is applied to yield four key results. First, analysis of the metabolic and physiological data shows evolutionary convergence in growth rate, glucose uptake rate, glycolytic ATP and NADH production but divergence in NADPH production strategies. Second, factorization-based analysis of the transcriptome revealed six conserved transcriptomic adaptations describing increased expression of ribosome and amino acid biosynthetic genes and decreased expression of stress response and structural genes. Third, correlation analysis identifies five tradeoffs underlying the transcriptomic profiles. Fourth, statistical tests leveraging ALE design identify four mutation-flux correlates and eight mutation-transcriptomic correlates that link mutations to systems level adaptation principles. Our total results reveal the dominant metabolic and regulatory constraints governing *E. coli* growth adaptation that either distinguish strains or are conserved principles.

## Introduction

Advancements in biotechnology have enabled the unprecedented detailing of microbial evolution. The process of evolution can now be studied in a controlled laboratory environment, where genome sequencing and phenotypic measurements are routine (Blount *et al.*, 2012; LaCroix *et al.*, 2015). Although studies utilizing genome sequences and fitness measurements have provided valuable insights ranging from the dynamics of evolution on long time-scales (Barrick *et al.*, 2009; Tenaillon *et al.*, 2016; Good *et al.*, 2017) to general features of epistasis (Kryazhimskiy *et al.*, 2014), evolutionary principles at the levels of gene regulation and metabolism remain unelucidated. Moreover, the generality of principles identified in experimental evolution studies is ambiguous since studies often focus on a single strain, not a species. For example, different strains of *E. coli* have been shown to exhibit diverse regulatory and metabolic functions and thus may have different constraints governing their evolutionary trajectories (Monk *et al.*, 2016). A fundamental multi-scale description of evolutionary landscapes may therefore be deduced through multi-omic measurements of different strain-specific experimental evolutions.

Towards revealing multi-scale features of evolutionary landscapes, researchers have taken transcriptomic and fluxomic measurements in their experimental evolution studies (Long *et al.*, 2018). However, it remains challenging to extract insights from these data types due to a lack of effective data analysis methods, especially for gene expression data sets. To date, no statistical correlation has been made between selected mutations and these multi-omics measurements. Our lab has recently shown the effectiveness of independent component analysis (ICA) to quantitatively interpret transcriptomic datasets in terms of transcription factors (Sastry *et al.*, 2019b). Therefore, ICA and novel statistical approaches may reveal fundamental regulatory principles and provide links between mutations and transcriptomic changes analogous to those seen in genetic association studies.

Here, we reveal multi-scale adaptation principles in the *E. coli* species by taking multi-omics measurements of six different strains throughout their adaptive evolution to glucose minimal media. Our total results reveal the dominant metabolic and regulatory constraints governing *E. coli* growth adaptation that either distinguish strains or are conserved principles.

## Results

### Consistent genetics in evolution of multiple *E. coli* strains

Six different *E. coli* wild-type strains exhibiting diverse genetics (K-12 MG1655, K-12 W3110, BL21, C, and Crooks) (**Fig. 1A**) were evolved to select for rapid growth. Independent triplicates of each strain were evolved under a strict selection pressure for growth in that the cultures never left the exponential phase under batch culture 37 C and M9 glucose (see **Methods**). Whole genome sequencing was performed for clones of all replicate lineages while 13-C fluxomics, RNA-seq, and physiological measurements were performed for a single replicate lineage (**Fig. 1B**). We find that all strains start with different growth rates but evolve to rates ranging between 0.98 and 1.11 hr^−1^ (D_t_ = 42 mins) (**Fig. 1C**). Some strains (W and Crooks) operate near this optimal in their wild-type state, while others require genetic mutations to achieve the observed optimal (MG1655, W3110, BL21, and C). We observed striking consistency in mutated genes, where each strain had at least one gene with a selected mutation in all replicate lineages (**Fig. 1D**). A total of seven genes (*pykF*, *zwf*, *spoT*, *mrdA*, *hns/tdk*, *rpoC*, *rpoB*) had selected mutations appear both in multiple strains and in more than one replicate lineage. The commonality of selected mutations indicated similar evolutionary constraints facing these strains and motivated inquiry of their metabolic and gene expression profiles.

**Figure 1:**
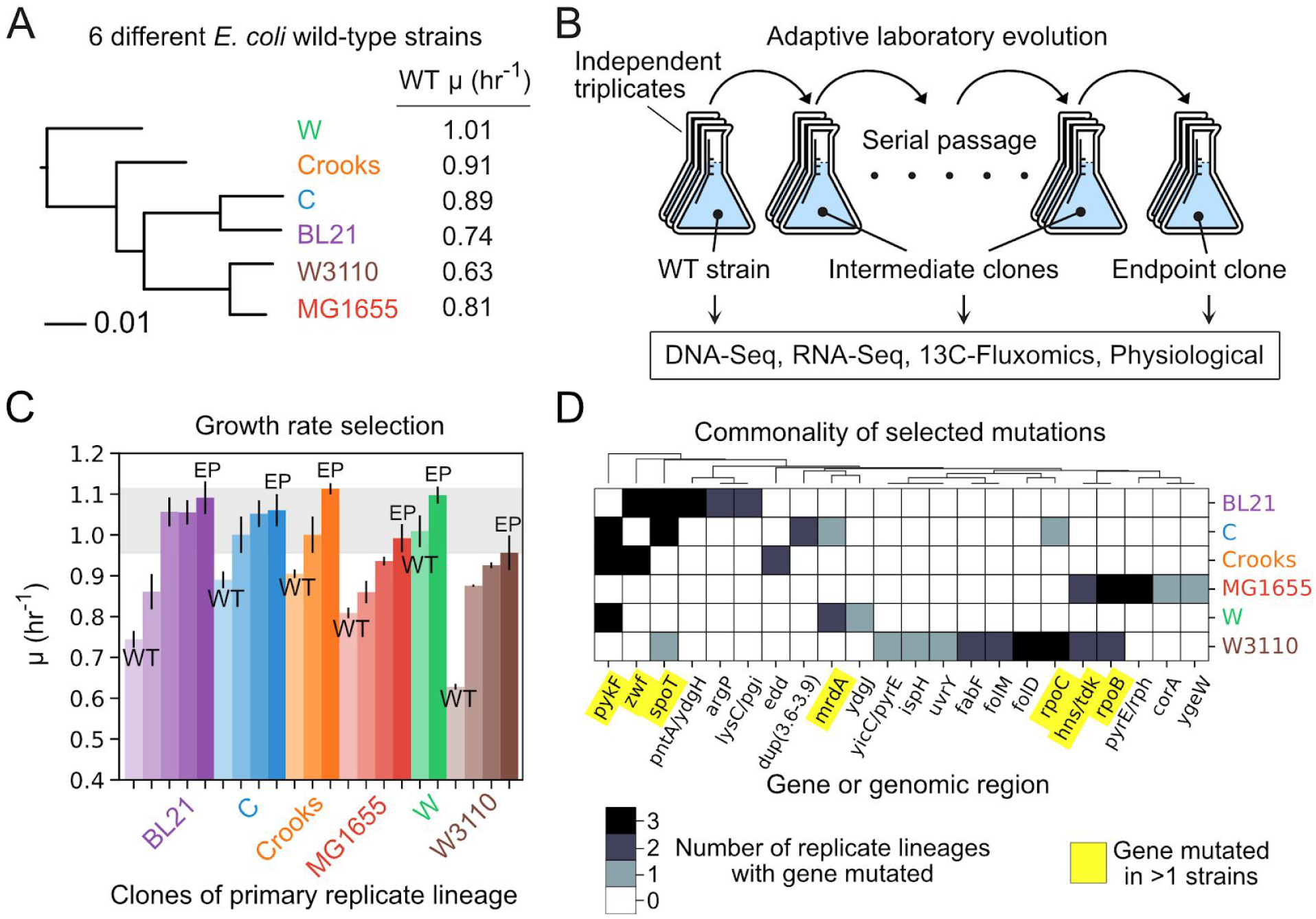
Overview of selected *E. coli* strains, experimental design, and key adaptation trends (**A**) Phylogenetic tree of six different *E. coli* wild-type starting strains utilized in this study. The wild-type (WT) growth rates (u) of the strains are noted. (**B**) Adaptive laboratory evolution was performed for each strain using independent triplicates. The wild-type (WT), evolved intermediate, and evolved end-point clones underwent multi-omics measurements. (**C**) Bar Plot of measured growth rates for wild type (WT), intermediate, and end point (EP) flasks for each strain. Clones are ordered left to right by trajectory. (**D**) Heatmap of gene-level mutation frequency across replicate lineages of each strain. The intergenic region between two genes is noted by a dash “/”.

### Characteristics of physiological and metabolic adaptations

Since a total of 8 selected mutations were in genes encoding metabolic enzymes—two of which appear multiple strains (*zwf*, *pykF*)—we hypothesized that the strains may be evolving towards similar metabolic states. We thus set out to examine convergent and divergent phenotypes along the ALE trajectory by performing statistical tests for each physiological and fluxomic measurement between the wild-type (WT) and end-point (EP) flasks for each strain (see **Methods**). Of the 187 total phenotypes, 64 were identified as convergent (i.e., points became closer together) and 6 were identified as divergent (i.e., points became further apart) with false discovery rate (FDR) less than 5% (**Fig. 2a**). Of the convergent phenotypes, we find that 86% (55/64) were growth-correlated (spearman rho<0.05, FDR<0.05) (see **Supplementary File 1**).

**Figure 2.**
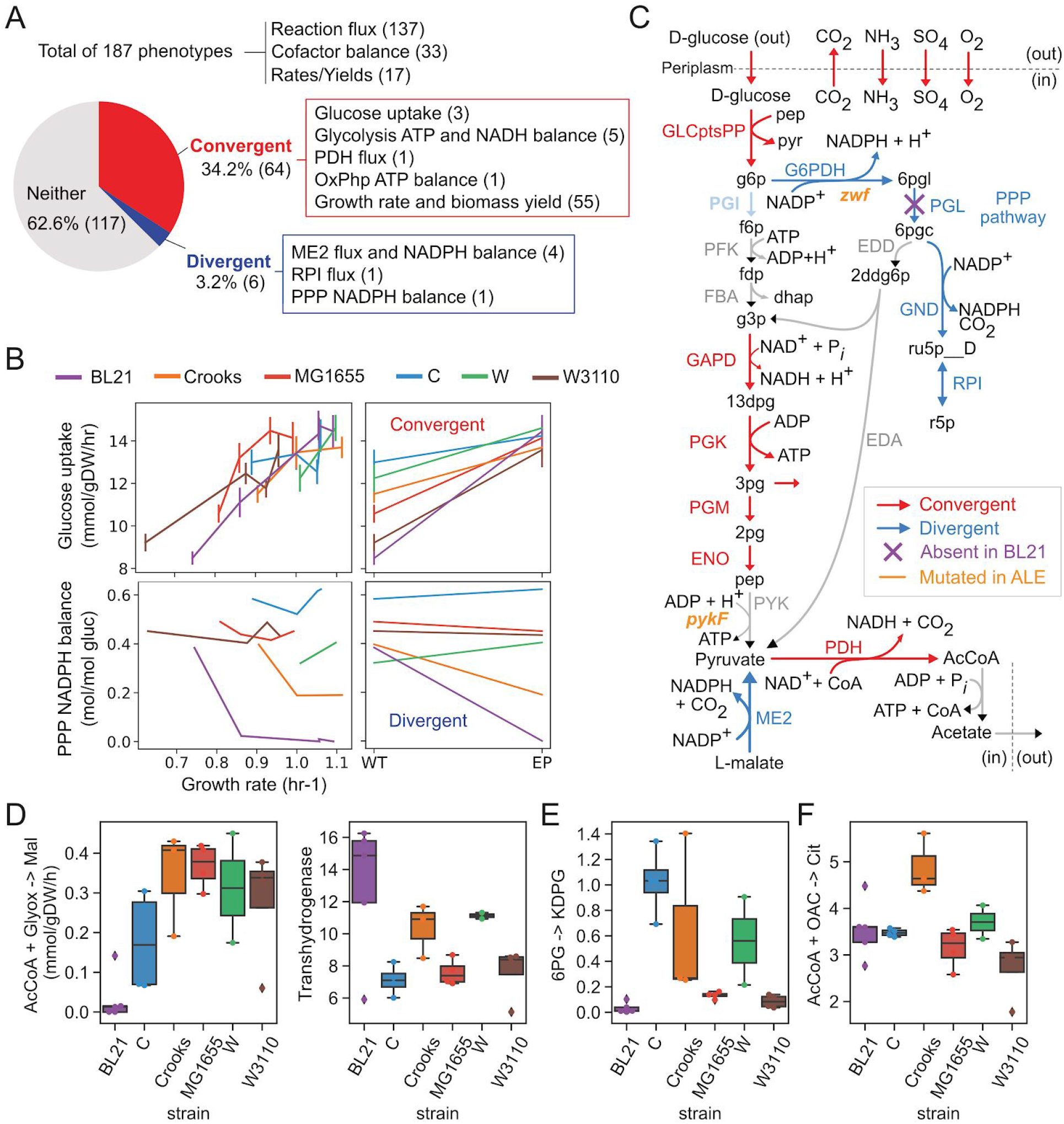
Adaptation in physiology and metabolism. (**A**) Pie chart describing the fraction of phenotypes that converge or diverge. Numbers in parentheses describe the number of related phenotypes. (**B**) Line plots of glucose uptake (top) and PPP NADPH balance (bottom) vs growth rate. Line plots and frequency distributions for WT and EP are plotted to the right for both cases. (**C**) Metabolic map of reactions in glycolysis, PPP, and exchange reactions colored according to whether they diverge or converge. Blue describes divergence and red describes convergence. (**D-F**) Bar plots of four reaction fluxes (absolute) that have strain-specific distributions. Abbreviations: abs, absolute flux (mmol/gDW/hr); rel, relative flux (mol/mol glucose); TCA, citric acid cycle; PPP, pentose phosphate pathway; ME2, malic enzyme; OxPhp, Oxidative phosphorylation; PDH, pyruvate dehydrogenase.

We find that the convergent features are related to glucose uptake, glycolysis and oxidative phosphorylation while the top ranked divergent features relate to NADPH production through malic enzyme (ME2) and pentose phosphate pathway (PPP) (**Fig. 2A**). Inspection of the ALE trajectories for the most convergent (Mann-Whitney U>169, *P*<5.7×10^−5^) and divergent (Mann-Whitney U=19, *P*=5.7×10^−5^) phenotypes showed that phenotypes do not monotonically increase/decrease along the ALE (i.e., not always increasing or decreasing along trajectory) (**Fig. 2B**). For example, although the glucose uptake rate has a significant net increase between WT and EP strains, four of the strains have one ALE jump where glucose uptake decreases. Principal component analysis of the fluxes showed that two components explain 93% of the variation and correspond to ATP production through oxidative phosphorylation and glycolysis (80%), and NADPH balance through pentose phosphate pathway and transhydrogenases (13%) (**Supplementary Figure 1**).

To determine whether specific reaction fluxes distinguish specific strains, we tested all fluxes for strain-specific distributions and found four subsystems specific to BL21, Crooks and C (ANOVA F-test, FDR<0.05). The BL21 strain uniquely had no flux through glyoxylate shunt while having the highest flux through transhydrogenase (**Fig. 2D**). Since BL21 can’t regenerate NADPH through PPP due to lacking the pgl gene encoding 6-phosphogluconolactonase (PGL) reaction activity (Meier, Jensen and Duus, 2012), the high transhydrogenase flux likely compensates to regenerate NADPH. Furthermore, we find that all BL21 flask lineages select for mutations in the intergenic region of a transhydrogenase (*pntA/ydgH*) (we test for mutation correlates later in this study) (**Fig. 1D**). C strain uniquely had high flux through the Entner-Doudoroff (ED) pathway while BL21, MG1655, and W3110 had almost none (**Fig. 2E**). Crooks uniquely had the highest flux through TCA (**Fig. 2F**). In total, these results describe convergent and divergent phenotypes that are either conserved or distinguish strains.

### Characteristics of transcriptome adaptation in *E. coli*

Underlying the phenotypic differences of these strains are differences in gene expression strategies. We thus set out to analyze the transcriptome of these strains by performing both differential expression analysis and a matrix factorization approach. Differential expression analysis showed that the number of differentially expressed genes (DEGs) generally decreases along the trajectory, with the exception of the last BL21 flask (**Fig. 3A**). To make sense of these expression changes, we applied an alternative RNA-seq analysis workflow that was shown to enable quantitative analysis of the *E. coli* transcriptome from the perspective of transcription factors (Sastry *et al.*, 2019b). The authors showed that independent component analysis (ICA) deconvolved a large compendium of *E. coli* MG1655 RNA-seq data into a linear combination of independent sources that reflect known regulons (“iModulons”), and source weightings (“iModulon activities”), which describe the global regulatory state (Sastry *et al.*, 2019a). Using the previous set of 92 iModulons, we transformed the flask-specific gene expression profiles into flask-specific iModulon activities (see **Methods, Supplementary Figure 2**).

**Figure 3.**
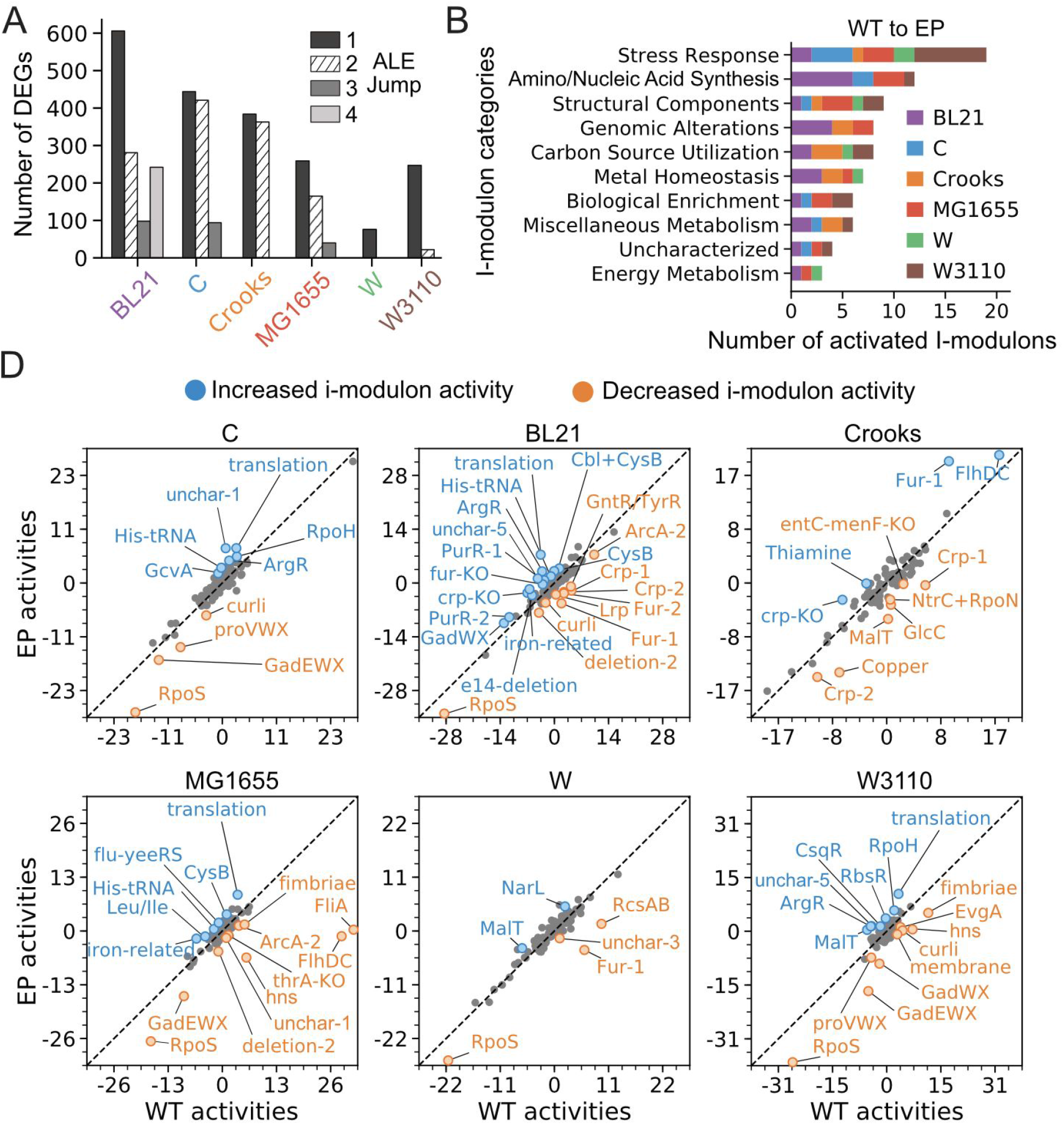
Characterization of gene expression adaptations. (**A**) Number of differentially expressed genes (DEG) for each strain-specific jump in growth rate during ALE. (**B**) Bar plot of total iModulon activation count in terms of iModulon functional category. The count is summed across the 6 strains activated ranked by the total number of times they were differentially activated between WT and EP flasks. (**C**) Bar plot of iModulons ranked by the total number of times they were differentially activated in an ALE jump. (**D**) Differential iModulon activity plots (DIMA). Comparison of iModulon activities between wild-type (WT) and end-point (EP) flasks for each strain. Significant altered iModulons are colored red and noted with text.

In order to first understand the different starting points of the strains, we tested for iModulons that distinguish WT expression profiles and identify a total of 38 iModulons (**Table 1**, FDR<0.005). For BL21, the iModulons imply an original environment that was cold (cspA), anaerobic and nitrate rich (ArcA-2), with gluconate (GntR/TyrR), allantoin, fructose, and arabinose (AllR/AraC/FucR) as possible carbon sources. For C, the identified iModulons hint at a background with high acidity and osmotic stress (EvgA, proVWX). The low OxyR activity in Crooks implies a WT environment facing low oxidative stress while high FliA activity in MG1655 implies that high motility was advantageous to its original environment. The relatively high GadEWX in W3110 implies an original environment with high acid stress.

**Table 1.** Strain-specific regulatory backgrounds. Table of conserved iModulons determined through ANCOVA with FDR<0.1. Bolded iModulons indicate that they also distinguish strains when looking at only WT flasks. iModulons with an Asterisk (*) describe iModulons distinguishing WT flasks but not all and double (**) describes iModulons specific to only EP flasks.

| Strain | iModulons distinguishing WT gene expression profiles |
| --- | --- |
| BL21 | AlIR/AraC/FucR, GntR/TyrR, CspA, deletion-1, ArcA-2, lipopolysaccharide, SrlR+GutM, YneJ, Ygbl, translation, membrane, gadWX-KO, fur-KO, RpoS, hns, sgrT, e14-deletion |
| C | NarL, flu-yeeRS, nitrate-related, uncharacterized-4, RbsR, NagC/TyrR |
| Crooks | OxyR |
| MG1655 | FliA |
| W | uncharacterized-1, ydcI-KO, ArcA-1, RcsAB, SoxS, Nac, CpxR |
| W3110 | PuuR, PurR-2, GadWX, curli, fimbriae, GadEWX |

To understand what iModulons changed the most throughout the ALEs, we performed differential activity analysis between the WT and EP flasks of each strain (see **Methods**). We find a total of 57 iModulons that were differentially activated at least once amongst the different strains (P<0.05, FC>2). The most commonly activated iModulons corresponded to stress response and amino/nucleic acid biosynthesis (**Fig. 3B**). The W3110 strain had the largest number of differentially activated stress response iModulons while BL21 had the most activated amino/nucleic acid biosynthesis iModulons. With respect to the total number of differentially activated iModulons, we find that BL21 has the most while W has the least (**Fig. 3D**), which reflects their respective change in growth rate. Of those activated, we find decreased activity in iModulons describing stress response (rpoS, gadEWX, rpoH, hns-related, proVWX) and motility (FlhDC, FliA, curli, fimbriae, RcsAB) while increased activity in iModulons describing translation machinery (translation), amino acid biosynthesis (ArgR, His-tRNA).

### Linear growth-dependent transcriptome adaptations conserved in *E. coli*

While differential activity analysis identifies general regulatory trends along the trajectory, it does not directly account for changes in quantitative growth rates or similarity between strains. We thus tested for iModulons that exhibit linear growth-dependence in all strains and identify six iModulons (**Fig. 4**, median Pearson |R|>0.75, median P-value<0.05). Of the six, three are positively correlated with growth-rate and describe the expression of ribosomal genes (translation), arginine biosynthetic genes (ArgR), and nutrient response (ppGpp). The other three iModulons are negatively correlated with growth-rate and describe stress response (RpoS, GadEWX) and structural assembly (curli). These results describe growth-dependent transcriptome adaptations that are mostly conserved in the *E. coli* species.

**Figure 4.**
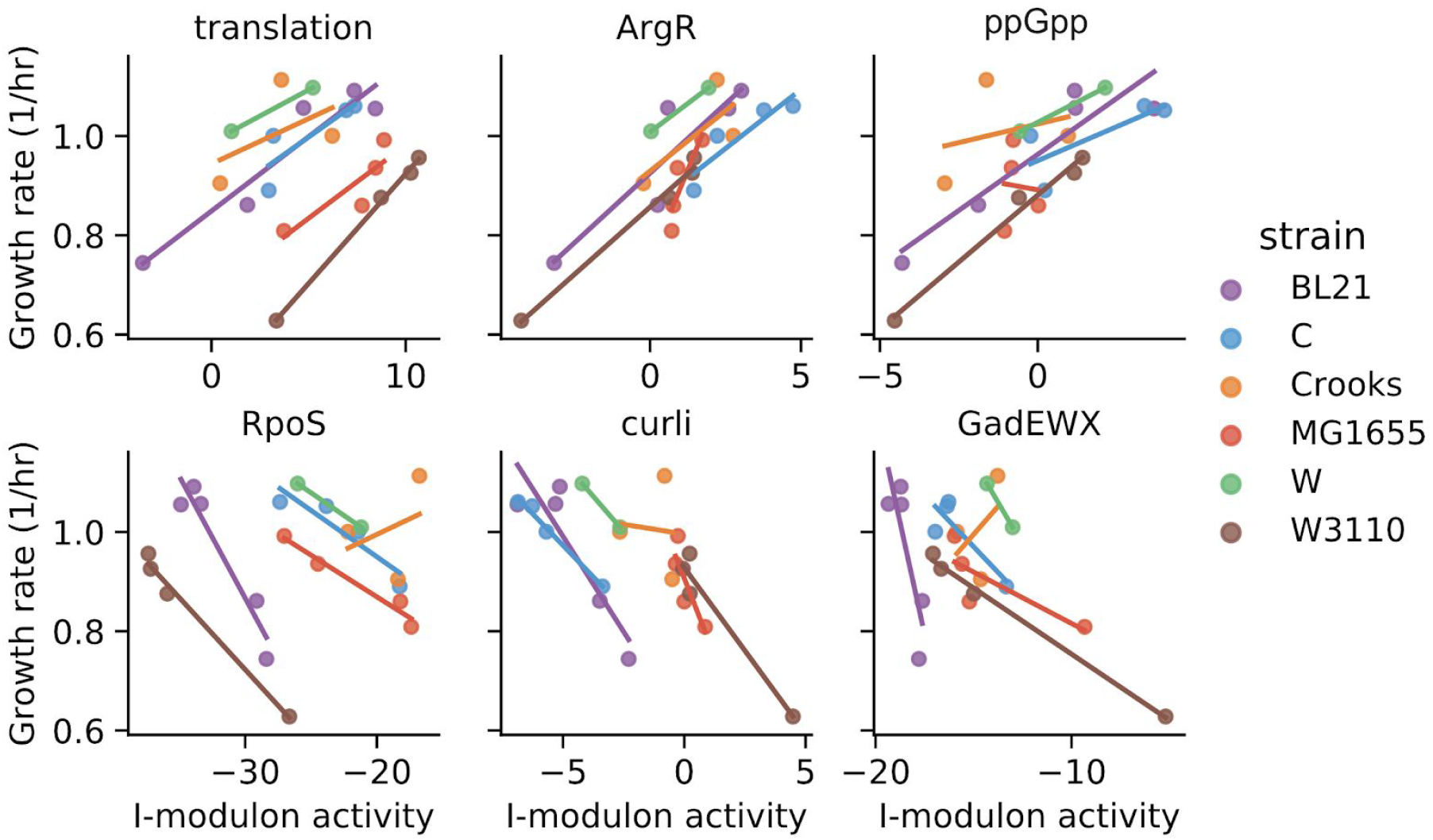
Conserved growth-dependent transcriptome. Strain-specific line plots of growth rate vs iModulon activity for six iModulons (median Pearson |R|>0.75, median P-value<0.05).

**Figure 5.**
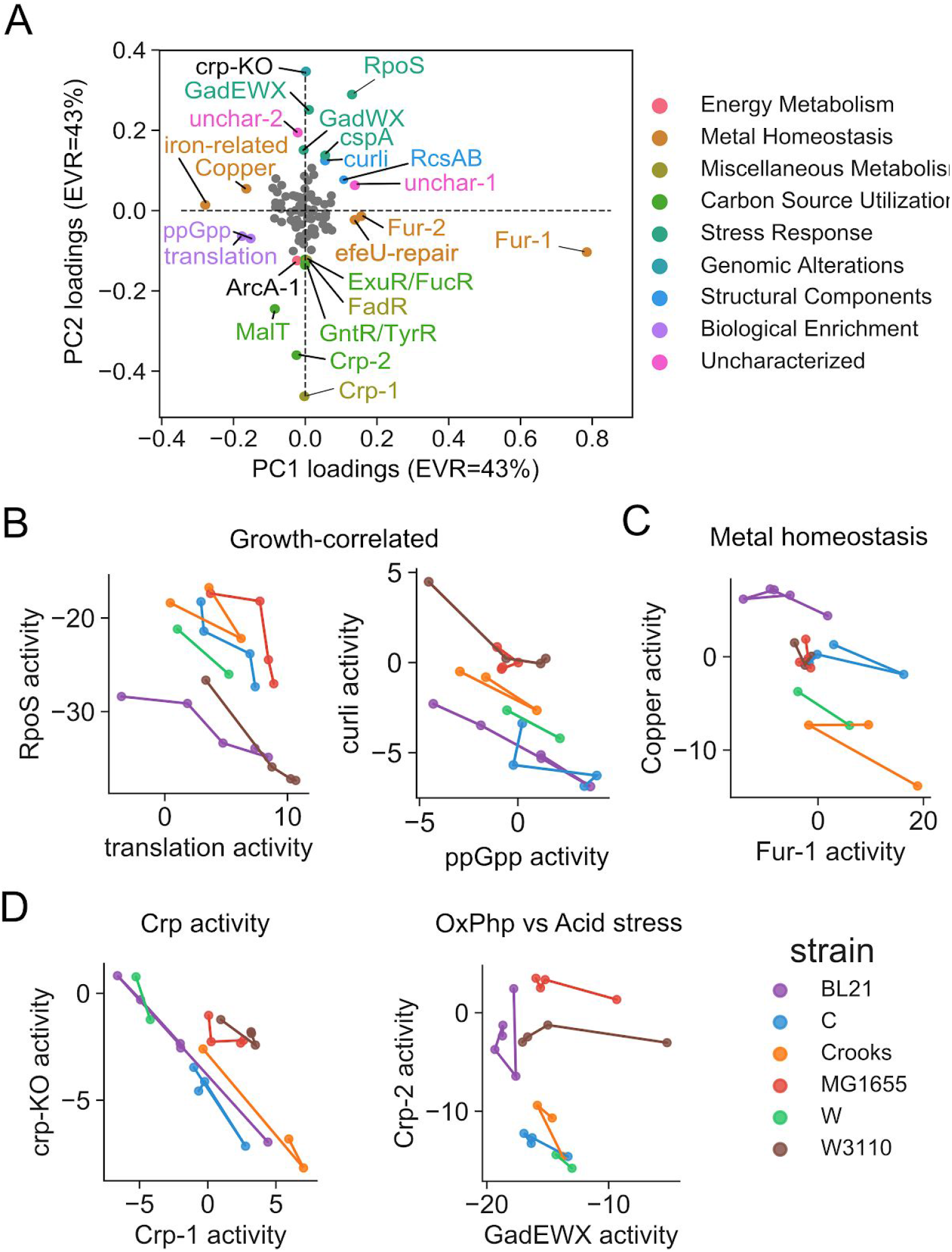
Regulatory trade-offs governing *E. coli* adaptations. (**A**) Plot of PCA loadings for components 1 and 2. (**B-D**) Strain-specific line plots for iModulon activities for trade-offs reflecting growth-correlated iModulons, metal homeostasis, crp activity, proton motive force.

### Regulatory trade-offs governing *E. coli* adaptation

The identification of both positively and negatively growth-correlated iModulons imply the existence of regulatory tradeoffs, and thereby a lower dimensionality of the iModulon activities (i.e., increased expression of certain genes requires decreased expression of others). We thus used principal component analysis to further decompose iModulon activities. Prior to PCA, we first transform the activity matrix (flask-specific) to the difference in flask activity along the trajectory (jump-specific) in order to identify components describing general adaptation trends as opposed to strain differences (see **Supplementary Figure XX** for PCA of flask-specific iModulon activities). We find that the first three PCA components explain the majority of the variance and have an explained variance ratio of 40%, 28%, and 12%, respectively. The first component describes activation of flagella machinery and is owed to the large deviation in FlhDC and FliA activity seen in the first MG1655 jump (**Supplementary Figure XX**). The second component describes metal-related iModulons (Fur-1, Fur-2, iron-related, efuR-repair, Copper) and growth-correlated iModulons (RpoS, translation, ppGpp). The third PCA component primarily describes carbon metabolism iModulons (Crp-1, Crp-2, MalT,) with positive weight and stress-response and structural iModulons with opposite weight (RpoS, GadWX, GadEWX, hns-related, CspA, curli). We test for negative correlations and identify a total of six potential tradeoffs (RpoS vs translation/ppGpp, Fur-2 vs translation/ppGpp, and Fur-1 vs Copper) in component 1 and (Crp-KO vs Crp-1, Crp-2).

### Statistical tests leveraging ALE design reveal key mutational effects

Comparing mutations is challenging due to the large number of genetic differences between strains. We therefore leveraged the directionality of the ALE data by transforming the flask-specific reaction fluxes and iModulon activities to jump-specific differences in flux and activity, thereby narrowing the view of genetic differences to those selected in ALE. Using the jump-oriented perspective of the data, we then tested for significant associations between jump-specific differences in reaction flux and iModulon activity with the selection of mutations at both the nucleotide and gene levels (i.e., gene level groups two different ALE mutations together if they appear in the same gene).

For the metabolic fluxes, we find four flux correlations primarily describing reactions involved in co-factor balancing (FDR<5%) (**Fig. 6A**). Specifically, *zwf* mutations are correlated with ΔG6PDH flux (NADPH balance through PP pathway), *pykF* mutations with ΔME2 flux (NADPH balance through Malic Enzyme), and *lysC* with ΔSUCCOAS flux (ATP and NADPH through TCA cycle). We find that the zwf mutation in Crooks is uniquely associated with ΔED pathway flux. For the iModulon activities, we identify eight mutation correlates that fall into four different iModulon functional categories describing stress response, motility, structural components, and carbohydrate metabolism (FDR<5%) (**Fig. 6B-E**).

**Figure 6.**
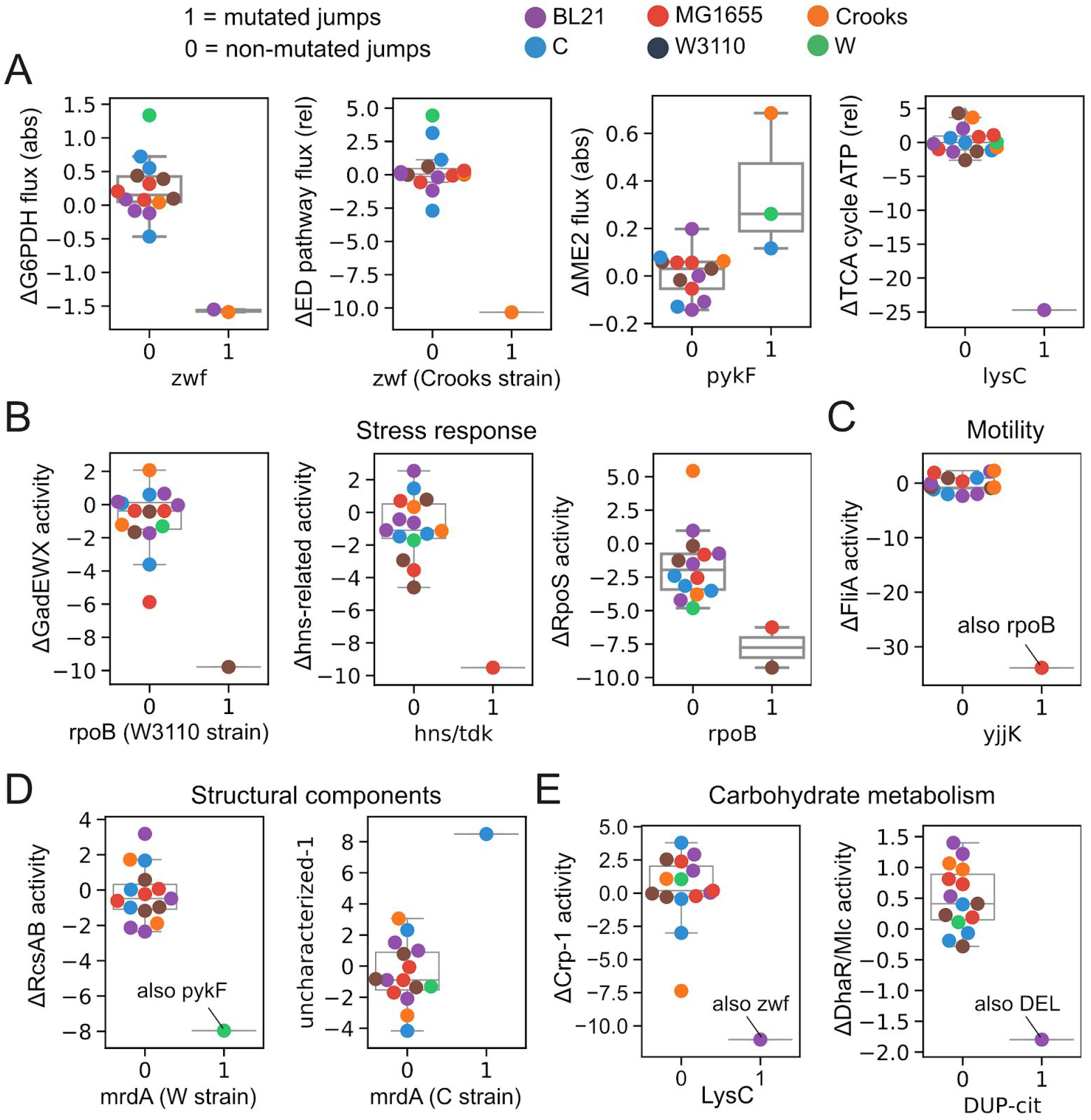
Mutation correlates. (**A**) Boxplots of significant correlations between mutations and changes in metabolic fluxes. The terms “abs” and “rel” in parenthesis refer to absolute flux (mmol/gDw/h) and relative flux (mol/mol gluc), respectively. (**B-E**) Boxplots of significant correlations between mutations and changes in iModulon activities. The boxplots are grouped by iModulon functional category. Genes with strains in parenthesis note a strain-specific mutation correlation. Mutations are grouped at the gene-level unless otherwise.

We find that similar statistical tests using DE fold changes instead of iModulon activities did not uncover any significant correlations. Notably, there are only 7 cases where the selection of a mutation coincided with significant differential expression of gene (**Supplementary Figure 2**). Factorization-based analysis therefore enables statistical associations between the transcriptome and selected mutations.

## Discussion

Taken together, our total analysis of the multi-strain ALEs revealed metabolic and transcriptomic adaptations principles of the *E. coli* species.

Characterization of the phenotypic data showed specific convergent and divergent features between the WT and EP flasks of these strains. It remains open how many peripheral phenotypes change with the core genes. Since the experimental condition was glucose minimal media, it remains unclear what principles are specific to glucose minimal media and which ones are not. Future studies may gain deeper insight by diversifying the measured phenotypes of these strains through high throughput approaches such as biolog plates.

Our ICA-based analysis of the transcriptome revealed key growth-correlated gene sets and tradeoffs governing *E. coli* adaptation.

By leveraging ALE design and the ICA-determined iModulon weights, we identified. Many of these associations make sense (i.e., *zwf* with ΔG6PDH flux, hns/tdk with Δhns-related iModulon activity) while others provide novel insights. Together, our results point to energy balance and proteome allocation (stress response, structural components, motility) as the dominant constraints governing *E. coli* adaptation. Including more samples would increase the identification of metabolic and regulatory features associated with mutations, providing a more clear picture of the logic underlying evolutionary selection.

Our results show that fluxomics and transcriptomics data types are valuable data types for characterizing adaptive landscapes.

## Methods

### Mann-Whitney U tests for identifying convergent and divergent phenotypes

To perform statistical tests for convergent and divergent features, we transformed the data vectors describing the mean physiological and fluxomics values for the size WT and EP flasks to vectors containing the pairwise distances amongst the points. The conversion resulted in a total of 15 points for each the WT and EP flasks. The transformation to pairwise distances accounts for how close the strains were at each point (i.e., convergence describes points coming closer together). Mann-Whitney U tests were then carried out to test whether the EP pairwise distances are smaller than the WT pairwise distances (i.e., the EP values are closer together than the WT values). We calculated the p-values using both a normal approximation implemented with the mannwhitneyu function in scipy stats and manual table acquired from from http://socr.ucla.edu/Applets.dir/WilcoxonRankSumTable.html. Both of the statistic estimates captured the general behavior, but the normal approximation was utilized due to the lack of table p-values for U statistics less than 36. We then selected the convergent and divergent features as those with a false discovery rate (FDR) less than 5% using the Benjamini Hochberg correction, implemented in the statsmodels package version 0.9.0 (Seabold and Perktold, 2010).

### Estimation of distinct growth groups

We determined an optimal clustering of measured growth rates using the k-means method implemented in the scikit-learn python package (Pedregosa *et al.*, 2011). The optimal number of total partitions was determined to be 7 (maximum silhouette score = 0.637). Furthermore, we performed linear regression using the python package seaborn [https://github.com/mwaskom/seaborn] to fit linear rate-yield lines for each distinct growth group. The k-means clustering of 7 growth partitions resulted in a set of best fit rate-yield lines (R^2^>0.85). We used the python packages matplotlib (Hunter, 2007) and pandas (McKinney, 2015) to perform visualization of growth groups.

### Differential expression analysis of RNA-seq

We performed differential expression analysis of the RNA-seq profiles between consecutive ALE flasks (i.e., ALE evolution stages) using the R package DESeq2 (Love, Huber and Anders, 2014). Specifically, differential expression was performed for each pair of flasks describing the before and after of an ALE experiment. We utilized an adaptive t prior shrinkage estimator (Zhu, Ibrahim and Love, 2018) to transform the log fold changes for better ranking and visualization of the differential expression results. Scatter plots of differential expression levels utilized the shrinked log fold changes (**Supplementary Fig. X**). We performed a sensitivity analysis of the p-value and Log_2_ fold change thresholds on determining sets of significantly expressed genes.

### Transcription factor and GO enrichments of differentially expressed gene sets

Transcription factors (TFs) and the genes they are known to regulate were obtained from a study (Fang *et al.*, 2017). GO terms were obtained from (Sastry *et al.*, 2019a). This list was used to perform hypergeometric enrichment analysis of the differentially expressed genes determined from RNAseq analysis. All differential expression analyses were compared to *E. coli* K-12 MG1655 as a reference. Therefore, the genes analyzed in this analysis were limited to the core set of 3306 genes contained in all strains. Genes between two samples representing the before and after of a specific ALEs that were significantly differentially expressed (p < 0.05, log2FC < > 2) where then used for hypergeometric enrichment. The enrichment scores for TFs and GO terms represent whether a significant number of differentially expressed genes are known to be regulated by a given TF or described by a specific GO term, respectively. The scipy (Jnes et al., 2001) stats package hypergeom was used to calculate hypergeometric enrichment values.

### iModulon analysis of RNA-seq data

In a previous study, we showed that application of independent component analysis (ICA) deconvolved a large compendium of *E. coli* MG1655 RNA-seq data into a linear combination of independent sources (“iModulons”), that reflect known regulons, and source weightings (“iModulon activities”), which describe the global regulatory state (Sastry *et al.*, 2019a). We therefore utilized the previous set of 92 iModulons (i.e., M from ICA(X)=MA) to deconvolve our multi-strain ALE expression data into iModulon activities. We utilized both the X (expression data) and S (iModulon network) matrices described in Anand et al (Sastry *et al.*, 2019a) to identify corresponding iModulon activities for our 46 RNA-seq. Specifically, the pseudoinverse was taken to compute the projection (see **Methods**, A=inv(M).dot(X)) (see **Supplementary Figure X**).

The iModulons Uncharacterized-6, Uncharacterized-3, and YYY were characterized as hns, CspA, and ZZZ respectively.

### Differential activity analysis of iModulons

Distribution of differences in iModulon activities between biological replicates were first calculated and a log-norm distribution was fit to the differences (Poudel *et al.*, 2020). In order to test statistical significance, absolute value of difference in activity level of each iModulon between the two samples were calculated. This difference in activity was compared to the log-normal distribution from above to get a p-value. Because differences and p-value for all iModulons were calculated, the p-value was further adjusted with Benjamini-Hochberg correction to account for multiple hypothesis testing problem. Only iModulons with change in activity levels greater than 5 were considered significant. Differential activity analysis was performed for all ALE jumps as well as between the WT and EP flask for each strain.

## Supplementary Discussion

### Cofactor balance convergence

Characterization of cofactor balance phenotypes showed convergence in the production and consumption of ATP via four subsystems: substrate uptake, biomass formation, glycolysis, and oxidative phosphorylation (**SI Fig. Xa**). We similarly observe convergence of these subsystems, in the production and consumption of NADH/FADH2, with the exception of substrate uptake (**SI Fig. Xb**). In contrast, we observe divergence in NADPH production/consumption strategies via the and ME2 (“other” corresponds to ME2) and convergence through biomass formation. We find that BL21 and Crooks strains underly the divergence of NADPH balance by decreasing PPP-based production of NADPH along their ALE trajectories (**SI Fig. Xc**).

### Transcription factor and GO enrichments of differentially expressed genes

To elucidate the DE gene sets, we performed transcription factor (TF) enrichments and found a total of 70 TFs enriched amongst the ALE jumps (FDR<0.1) (see **Methods**, **03_supplementary_file_DE.xlsx)**. Of these TFs, the most frequently enriched were *fur*, *arcA*, *Sigma70*, *cra*, *iscR*, *ryhB*, and histidine (Number of jumps > 4) (**Fig 3b**). We similarly performed GO enrichments and found that the most enriched GO terms (FDR<0.1) corresponded to respiration, ribosomal, TCA cycle, enterobactin, and histidine biosynthesis **Fig 3b**), which reflect the enriched TFs (see **SI Table 3**). The GO term ‘iron-sulfur assembly’ is enriched in the last ALE jump of BL21 and MG1655, in which an increase in DE genes is observed. In total, the DE genes generally point to respiratory control as the dominant strategy in this ALE.

### iModulons

Together, the 92 iModulons explained 52% of the expression variance of the multi-strain core genome, where they explained the most expression for MG1655 (67.78%) and the least for C (44.23%) (**Supp. Fig. 3B**). The specific iModulons explaining the largest percentage of the expression were described by *rpoS* general stress response iModulon, *gadEWX* acid stress response, and flagella motility (FliA, FlhDC) (**Supp. Fig. 3B**).

### Comparison of iModulons and proteome sectors

Comparing the 65 iModulons to the 6 proteome sectors, we found that the 65 iModulons controlled 923 genes that describe on average 26% of total TPM in each sample, in contrast to the 75% described by the 905 proteome sector genes (see **Fig. Xb**) (see **Methods**, thresholding the iModulon gene weights to generate a binarized M matrix). Only 32.4% (293/905) of the proteome sector genes were controlled by the iModulons. The iModulon genes generally did not line up well with the proteome sectors, with the most overlap occurring in proteome sector C with 58% of genes (53/91) (see **Supplementary figures/00/sectors_imods_CUTOFFS-0.csv**).

## Supplementary Figures

**Supplementary Figure 1.**
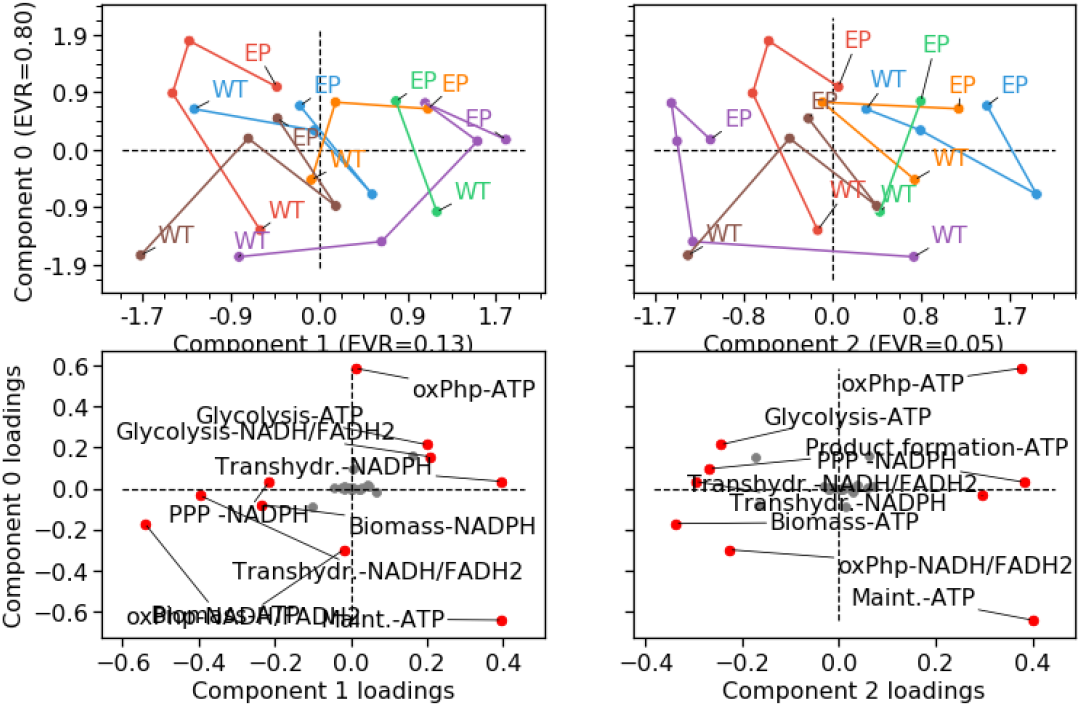
Principal component analysis of fluxomic data.

**Supplementary Figure 2.**
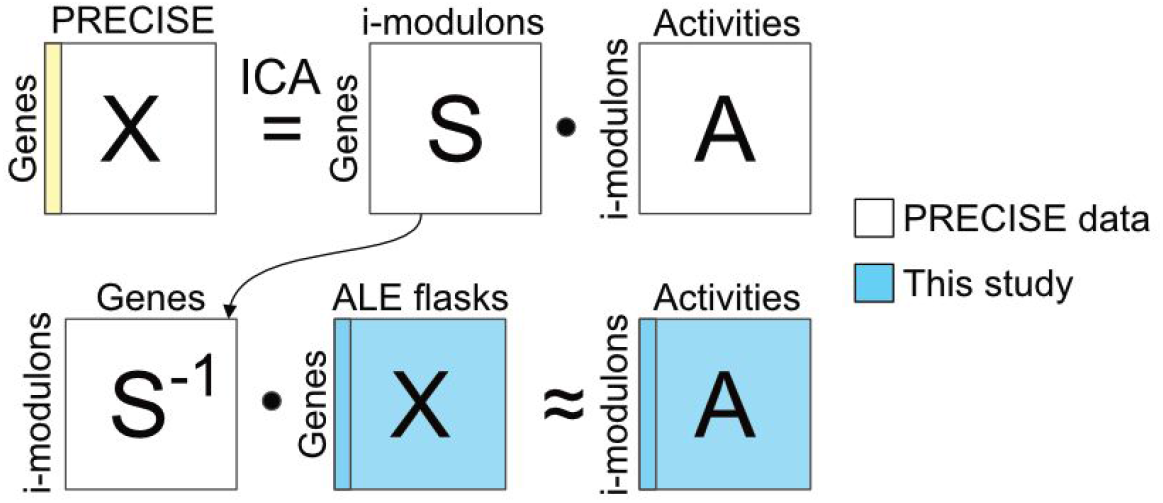
Schematic of ICA-based approach. The S matrix determined by the PRECISE database was utilized to estimate the iModulon activities describing the samples in our dataset.

**Supplementary Figure 3.**
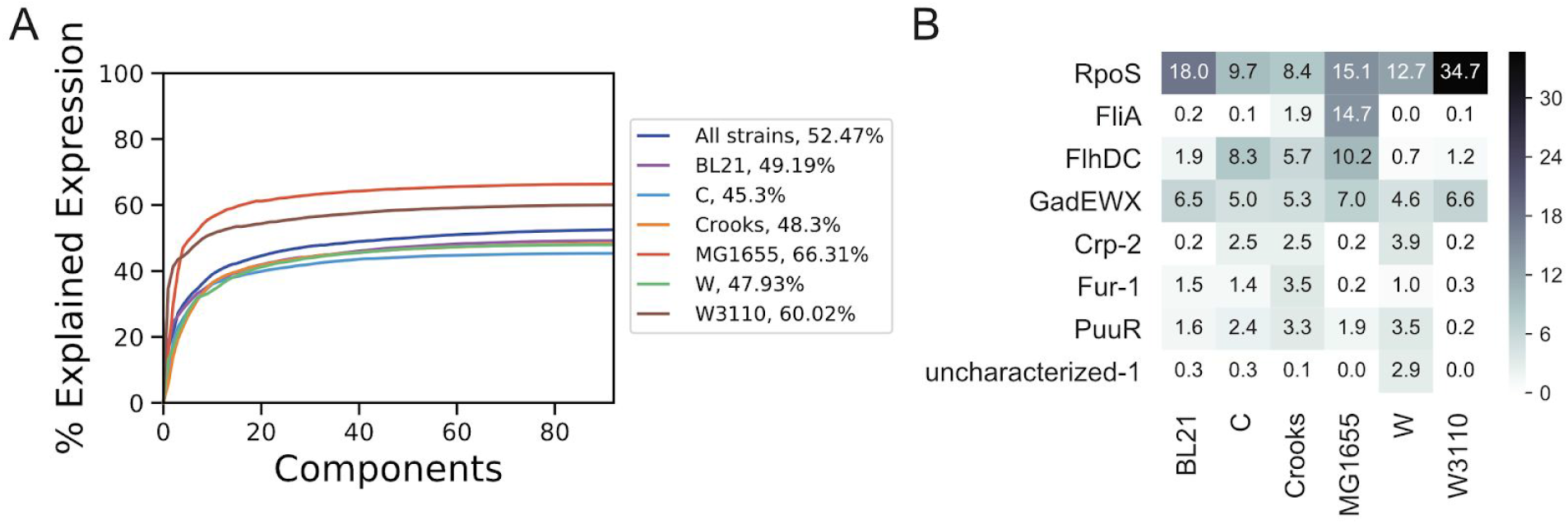
Explained expression by PRECISE iModulons. (**A**) Explained expression by mapped *E. coli* iModulons taken from Sastry et al. (**B**) Heatmap of percent explained expression per iModulon across the 6 strains. Only 8 iModulons are shown that explain more than 3.5% across the strains are shown and vertically ordered from most to least.

**Supplementary Figure 4.**
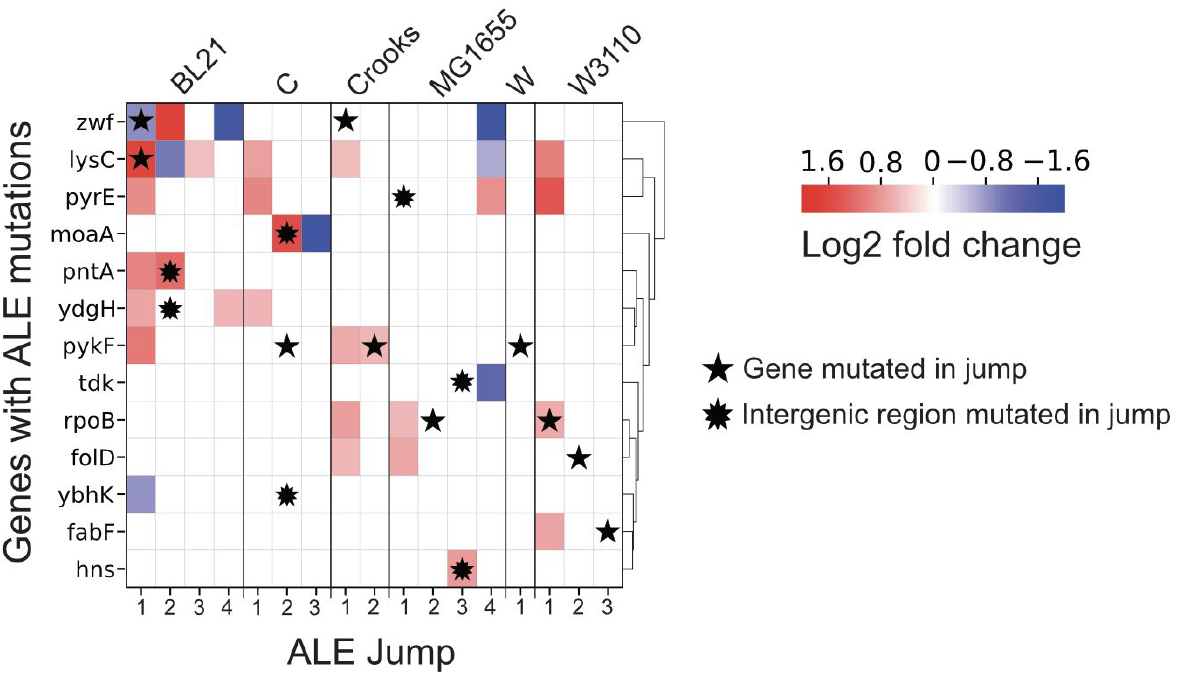
Clustered heatmap of expression changes for genes containing at least one mutant across all strain-specific ALE jumps. Log2 fold changes are only shown for significant differentially expressed genes in each jump.

## Code availability

Code is available upon request.

## Supplementary Files

Supplementary

## Notes

**Source of support** This research was supported by Novo Nordisk Foundation (NNF10CC1016517).

### Competing Interest Statement

The authors have declared no competing interest.

